# Inflammatory Proteins are Highly Variable During a Natural Stressor: A Call for Intensive Longitudinal Blood Microsampling

**DOI:** 10.64898/2026.01.09.698715

**Authors:** Daniel P. Moriarity, Andrea C.M. Miller, Durga D. Thota, Aaron Miller, Kevin Trent, Tory Eisenlohr-Moul, Thomas McDade, Michael P. Snyder, George M. Slavich

## Abstract

Much research investigates how inflammation might play a role in stress-mediated diseases such as heart disease and depression. However, standard longitudinal designs often have measurement lags of months or years between assessments. This status quo is at odds with evidence that inflammatory biology and stress are both highly dynamic and would require much shorter time lags to quantify change as it naturally occurs, with minimal risk of confounding or effects diluting over time. Inspired by the use of intensive longitudinal data collection in psychological research, this study collected high frequency (every 3 days) blood microsamples over 22 days as 86 incoming college students transitioned onto campus (total observations = 622 blood samples). Samples were analyzed for CRP, TNF-α, IFN-γ, IL-1β, IL-2, IL-4, IL-6, IL-10, IL-12p70, and IL-17A. Results using both bivariate correlation and intra-class correlation coefficients (which can be used for future power analyses for longitudinal research with these proteins) demonstrate low reliability. Notably, hierarchical linear models demonstrated that, even at this frequency of assessment, most proteins were characterized by less than 6% stable between-person differences. These results provide compelling evidence that investment in intensive longitudinal data collection of immune proteins is critical for understanding how inflammatory biology might function in psychosocial models of risk and resilience.

## Introduction

The field of psychoneuroimmunology has provided compelling data about the existence of bidirectional interplay between immune functioning, neurobiology, and thoughts, feelings, and behaviors. Much of this research focuses on the role of psychosocial stress in modulating the immune system^1–10^, as well as the psychological risk and resilience factors that influence this path (e.g., emotion regulation^11–14^), and the associated health consequences (^1,6,15–17^). However, common longitudinal study designs fail to answer a fundamental question: How reliable are inflammatory proteins over periods of naturalistic stress?

Understanding the reliability of a system is imperative to understanding how frequently it needs to be assessed to maximize the theoretical and translational value of research. The less temporally reliable a variable is, the more often it needs to be assessed for data to accurately reflect changes in the system as they occur—a critical feature to both understand (a) what drives changes in the variable and (b) how changes in one variable might influence changes in others. Conversely, more constant variables are generally able to be assessed less often (perhaps, only once for highly stable traits depending on the intended use) without fear of missing theoretically or clinically meaningful change. By mapping assessment frequency onto the natural dynamics of the systems of interest, longitudinal studies are better able to capture interactions between variables as they naturally unfold, which is essential to refining etiological theories about stress, immune functioning, and health outcomes^18^. Otherwise, effect sizes are more susceptible to confounding (when measurement lags are overly long) or downward bias (when measurement lags are too short for processes to unfold). Importantly, methodologists recommend that until ideal assessment frequencies are known, scientists should prefer to risk sampling more frequently than necessary than risk under-sampling^19^. This is both because over-sampling (as long as the total duration is long enough) will still capture change as it occurs (albeit less efficiently that ideally sampled data) and the resulting data can facilitate analyses to determine the ideal frequency.

This issue is not specific to psychoneuroimmunology; in fact, it is pervasive throughout psychological research^20,21^, partially inspiring the rapid adaptation of intensive longitudinal research designs (including ambulatory assessments) in psychology. The sensitivity of psychoneuroimmunological associations to small differences in measurement lags are driven home by (a) a study that found that negative affect (assessed 5x/day for 14 days leading up to a blood draw) was more strongly correlated with a cytokine composite (sampled from blood) when the affect assessment was closer to the blood draw^22^ and (b) a recent case study finding differential associations between well-being and urinary TH1 immune activation based on measurement lag when both were assessed every 12 hours for 63 days^23^.

Investigations into the temporal physiometrics^24–26^ of inflammatory biology have highlighted lack of reliability in inflammatory proteins over the months-to-years long time lags often observed in published research (e.g., 18-month reliability in salivary inflammatory proteins^27^) and even within the same day (i.e., diurnal variations in salivary C-reactive protein^28^, 3 hour differences in salivary IL-8 and IL-1β^27^). It is plausible that inflammatory proteins are even less reliable during prolonged periods of stress and adjustment given that stress itself (as indexed by the Perceived Stress Scale), a known driver of inflammatory upregulation^1,7,9^, does not hold static over prolonged periods of time^29^.

Despite the immense investment in and theoretical importance of stress research in psychoneuroimmunology, no studies have tested the reliability of blood-based inflammatory proteins during periods of naturalistic stress when psychosocially induced immune reactions (and their acute sequalae) are most likely to occur. Additionally, initial physiometric investigations have either focused on change across a single day or change over measurement lags of months-to-years. To complement the known dynamics of both psychosocial stress and inflammatory proteins, it is imperative to quantify the temporal reliability of peripheral immunology using intensive longitudinal data across a period of stress and adjustment^18^.

### The Present Study

To address this issue, we used data from Project MHISS^30^, which recruited incoming 1^st^ year students to complete a novel intensive longitudinal protocol during the transition to campus which lasted 22 days. Blood microsamples were collected every 3 days to assess for trajectories of inflammatory proteins. The transition to college was selected as an ecologically valid stressor because it is a high intensity social stressor that combines major interpersonal, lifestyle, and goal-oriented adjustments—increasing the likelihood of meaningful fluctuations in stress (and consequently stress biology) during the study period. Truly, for most college students, it is the largest developmental transition they have ever experienced. The present report characterizes the temporal reliability of C-reactive protein (CRP), tumor necrosis factor (TNF)-α, interleukin (IL)-6, IL-4, IL-2, IL-1β, IL-17A, IL-12p70, IL-10, and interferon (IFN)-γ to provide fundamental information about the reliability of these proteins, at this measurement frequency, during an intensive naturalistic stressor and evaluate the necessity of intensive longitudinal immunological data collection in observational stress research.

Reliability will be quantified both using (a) average Pearson correlation coefficients between two time points separated by three days and (b) intraclass correlation coefficients (ICCs) that compare the degree of within-person variability (i.e., the degree of change over time in each individual’s proteins) and between-person variability (i.e., protein levels that are stable across the study). These metrics were chosen because (a) they quantify complementary, yet distinct, facets of temporal reliability, (b) they are relevant for different study designs (two timepoint vs. two or more timepoints, respectively), and (c) are both useful estimates for power analyses for popular longitudinal statistical methods (e.g., linear regression predicting future inflammation covarying for baseline inflammation and hierarchical linear models, respectively).

## Method

### Participants

Participants were recruited from a large public university in California, USA, via electronic screeners delivered by the Office of Admissions. To be eligible, participants had to: (a) be an incoming first-year undergraduate student, (b) be 17-19 years old (to avoid recruiting participants who had a greater likelihood of having a major life transition before the move to college such as military service or moving for a job or be at a different state of social development/salience), (c) be moving at least 100 miles into a campus dormitory (to avoid social stress buffering via proximity to family), (d) not have chosen their own roommate, and (e) be fluent in English. In addition, participants needed to be comfortable drawing their own blood microsamples and not have current severe immunological diseases or be using strong immune-modulating drugs. Descriptive statistics of demographics variables for the entire analytic sample (86 individuals) are presented in Tables 1 and 2.

**Table 1.** Demographics– Continuous variables.

| Characteristic | Range | Mean | Median | SD |
| --- | --- | --- | --- | --- |
| Age | 17-19 years | 18.2 years | 18.0 years | 0.40 years |
| Subjective SES | 1-10 | 6.3 | 6.0 | 1.8 |

Note: SD = Standard deviation, SES = Socioeconomic status.

**Table 2.** Demographics Data – Categorical variables.

| Characteristic | Number (%) |
| --- | --- |
| **Sex at birth** |  |
| Female | 66 (76.7%) |
| Male | 20 (23.3%) |
| **Gender (Most strongly identified)** |  |
| Genderfluid | 1 (1.2%) |
| Gender non-binary | 1 (1.2%) |
| Man | 20 (23.3%) |
| Two spirit | 1 (1.2%) |
| Woman | 59 (68.6%) |
| Not listed or prefer to self-describe | 1 (1.2%) |
| I feel that multiple identities better describe the way in which I identify | 3 (3.5%) |
| **Sexuality (Most strongly identified)** |  |
| Bisexual | 12 (14.0%) |
| Heterosexual | 48 (55.8%) |
| Lesbian or gay | 5 (5.8%) |
| Mostly heterosexual | 7 (8.1%) |
| Queer | 5 (5.8%) |
| Questioning | 1 (1.2%) |
| Pansexual | 1 (1.2%) |
| Demisexual | 1 (1.2%) |
| I feel that multiple identities better describe the way in which I identify | 2 (2.3%) |
| I do not use a label | 4 (4.7%) |
| **Race** |  |
| Black or African American | 6 (7.0%) |
| East Asian | 20 (23.3%) |
| Hispanic or Latina/e/o/x | 19 (22.1%) |
| Native Hawaiian or Pacific Islander | 3 (3.5%) |
| South Asian | 8 (9.3%) |
| White | 40 (46.5%) |
| Not listed or prefer to self-describe | 3 (3.5%) |
| Prefer not to say | 2 (2.3%) |
| **Family Income** |  |
| More than \$200,000 | 23 (26.7%) |
| \$100,000-\$200,000 | 20 (23.2%) |
| \$50,000-\$100,000 | 9 (10.5%) |
| \$25,000-\$50,000 | 15 (17.4%) |
| Less than \$25,000 | 6 (7.0%) |
| Prefer not to say | 13 (15.1%) |

Note: Percentiles for category might sum to >100% due to (a) rounding and/or (b) the option for participants to select all racial identities that apply.

### Procedures

Blood microsamples were collected on Days 1, 4, 7 (1^st^ full day on campus), 10, 13, 16, 19, and 22. Participants who missed more than 2 blood draws were excluded from the rest of the study. Participants were instructed to complete the blood draws prior to their first meal/caffeine intake and between 8:00 am and 11:00 am local time to control for diurnal fluctuations^28,31^. The timing and quality of the blood draw collection (e.g., that the adhesive was removed so that the sample could dry) were verified by uploading a timestamped image of the used device and its box indicating the participant ID number and blood sample number. If participants reported a fever or felt ill, they were instructed to not sample blood. Samples were dried within the device and shipped overnight to a laboratory at Stanford University, where they were labeled, processed, frozen, and stored at -80^°^C before analysis.

Samples were collected via TASSO M-20 devices, which were distributed to participants along with an instructional video, written guidelines, and an FAQ sheet to assist with their use. Research staff were available during study hours to video chat with participants to successfully execute blood microsampling. TASSO M-20 devices attach to the shoulder with an adhesive, and when the button is pressed, a vacuum forms and a lancet pricks the surface of the skin to collect dried whole capillary blood. The vacuum draws ∼80 μL blood out of the capillaries into a container attached to the bottom of the device. The Tasso-M20 collects 4 samples of 17.5μL each with a coefficient of variation (CV) of <5%. Tasso-M20’s dried blood samples can collect and stabilize samples for drugs that are small molecules, proteins, antibodies or contain nucleic acids.

Because some students lived in different time zones prior to moving to campus, the analyses for this project are restricted to individuals who did not cross time zones during their move to campus (which would result in disconnect between circadian rhythms and local time, as well as result in systematic differences in the number of hours between assessments). Two individuals were removed (16 observations) due to use of immune-modulating medications, and 45 observations were removed due to sample collection issues (i.e., failure to remove adhesive that allows blood to dry, overfilled plugs). This resulted in an analytic sample size of 86 individuals and 622 blood samples. To ensure that results of the intra-class correlation coefficients were not influenced by differences in how many blood draws were completed by each person (because more datapoints generally leads to more reliable estimates), the ICCs calculated below only include individuals with data for each respective immune protein at all 8 occasions (specific number of participants and observations for each protein are reported).

## Measures

### Immune Proteins

Inflammatory proteins were analyzed at Northwestern University. Samples were assayed for C-reactive protein (CRP) and a 9 protein, ultrasensitive multiplex (the MSD Proinflammatory (human) Panel 1 Kit), which included interferon (IFN)-γ, interleukin (IL)-1β, IL-2, IL-4, IL-6, IL-10, IL-12p70, IL-17A, and tumor necrosis factor (TNF)-α. All proteins were assayed in a single batch. The Tasso samples we analyzed were eluted overnight in 218.0 μL of an assay buffer (phosphate buffered saline, 0.1% Tween-20) at 4C. A modified protocol for Tasso samples was used, resulting in 50μL of each sample added to the plate in triplicate. After samples were added the remaining procedures followed the kit protocol (MSD #K15396S). At the end of the assay procedure the plate was read using a Meso Scale Discovery QuickPlex SQ 120 Imager, with data being processed in the Discovery Workbench 4.0 Analysis software. C-reactive protein was analyzed using an in-house ELISA assay originally designed for dried blood spot analysis^32^ and modified for use with Tasso M20 samples. Samples were eluted the same as in the Multiplex assay, further diluted, and 100μL of diluted eluate was added to each assay plate in duplicate.

CRP was run in duplicate. All multiplex assays were run in triplicate. Intra-assay coefficients of variation (CVs) were calculated for each immune protein as a measure of reliability. The use of triplicate cytokine was to counteract known issues with reliability in multiplex assays in two ways. First, averaging across three runs will typically provide more reliable estimates than two^33^. Second, it allowed for identification and removal of outliers within the triplicates. To do so, each triplicate value was compared to the average of the other two triplicates for a given sample. When one triplicate was at least 5x the size of the average of the other two, it was removed and the average + CV for that sample was recalculated. Note that this facilitates cleaning likely invalid data points prior to aggregation while preserving the total number of observations for modeling. The intra-assay coefficients of variation (CVs) and number of triplicate level outliers are available in Table 3.

**Table 3.** List of analytes, coefficients of variation, and triplicate level outlier rate.

| Analyte | Median Intra-assay<br>Coefficient of Variation | Triplicate-level outliers<br>(frequency/percentage) |
| --- | --- | --- |
| CRP | 1.8% | N/A |
| IFN-γ | 9.2% | 5 (0.8%) |
| IL-1β | 1.5% | 7 (1.1%) |
| IL-2 | 29.2% | 14 (2.2%) |
| IL-4 | 8.7% | 6 (1.0%) |
| IL-6 | 3.8% | 3 (0.5%) |
| IL-10 | 11.8% | 12 (1.9%) |
| IL-12p70 | 8.9% | 26 (4.2%) |
| IL-17A | 12.4% | 6 (1.0%) |
| TNF-α | 10.5% | 10 (1.6%) |

Note: CRP = C-reaction protein; IFN = interferon; IL = interleukin; TNF = tumor necrosis factor

### Statistical Analysis

The analytic plan for this study was pre-registered https://osf.io/5qy6h/overview?view_only=a55e0138892747e29de6b19c4691b2ab. The following deviations were made from the pre-registration. First, bivariate correlations were calculated as an additional index given their relative familiarity and ease-of-interpretation relative to intra-class correlation coefficients and their relevance to data with only two time points. Second, we incorporated the ICCs into Spearman-Brown Prophecy Formula to better put these psychometric results into context for study design. Third, we correlated the ICCs with intra-assay CVs to evaluate the potential for assay reliability to influence results.

Correlations were calculated using the himsc package^34^. Correlations for each 3-day measurement lag were averaged to determine the average 3-day retest reliability of each protein. Hierarchical linear models were estimated using lme4 package^35^. Models specified each protein as the outcome (i.e., 10 different models were estimated) with the only predictor being the random intercept (aka an “empty” model). Intra-class correlation coefficients (ICCs) were used to estimate the reliability of each protein using the formula *ICC* = *BetweenVariability*/(*BetweenVariability* + *WithinVariability*). Briefly, ICCs are the ratio of stable between-person variability (indexed by the intercept) of each protein divided by the total variability. As such, higher ICCs that approach 1.00 are indicative of highly reliable proteins with which between-person comparisons would accurately represent the majority of the variability. Conversely, lower ICCs closer to 0 are indicative of highly dynamic proteins with higher within-person variability. To maximize interpretability, ICCs are discussed in terms of within-person variability (calculated by converting the ICC to a percentile representing the proportion of variability attributable to reliable, between-person differences and subtracting that from 100%) below. To test whether reliability estimates were associated with potential measurement concerns of the immune assays, ICCs were correlated with the intra-assay CVs for each protein using Pearson correlations.

To evaluate the sensitivity of these stabilities to slightly longer durations between assessments, secondary models will be re-estimated using every other blood sample (i.e., on days 1, 7, 13, and 19). One of these secondary models (specifically, predicting IL-1B) was singular, precluding calculation of an ICC due to variability due to the intercept being negligible. In this situation, the model was re-estimated with weakly informed priors in the *brms* package ^36^, which resulted in non-singular fit.

More assessments will generally lead to higher ICCs. As such, some researchers might be interested in understanding how many assessments would be necessary to have a “reliable” assessment of between-person differences over a period of time. The Spearman-Brown Prophecy Formula was used to determine the number of blood samples that, based on the reliability observed using the 3-day lag, would be necessary to achieve an ICC of .5 (i.e., the estimate is half between-person variability, half within-person variability). This threshold was selected for illustrative purposes as it is a recommended threshold for moderate reliability^37^. Analytic code is available at https://osf.io/5qy6h/overview?view_only=a55e0138892747e29de6b19c4691b2ab, which can be modified for other ICCs of interest to the reader.

## Results

### Bivariate Correlations Between Timepoints

Average 3-day retest reliability for each protein (in descending order) was: IL10 (*r* = .70), CRP (*r* = .60), IL4 (*r* = .50), IL17-A (*r* = .46), IL2 (*r* = .42), IL12p70 (*r* = .35), TNF-α (*r* = .31), IL1β (*r* = .30), IL6 (*r* = .20), IFNγ (*r* = .03).

### Within vs. Between-person Variance

After removing participants who did not have data at all eight observations the number of people and observations for each protein were: CRP = 240 observations (30 people), IFNγ = 280 observations (35 people), IL10 = 248 observations (31 people), IL12p70 = 160 observations (20 people), IL17A = 272 observations (34 people), IL1B = 264 observations (33 people), IL2 = 176 observations (23 people), IL4 = 272 observations (34 people), IL6 = 280 observations (35 people), TNFα = 280 observations (35 people). In order of most reliable to least reliable (see Table 4 for numeric results, see Figure 1 for Riverplots that visualize the disaggregation of between-vs. within-person variability for each protein): CRP had 54.18% within-person variability, IL-10 had 79.26% within-person variability, IL-4 had 94.96% within-person variability, IFN-γ had 96.11% within-person variability, IL-17A had 98.07% within-person variability, IL-2 had 98.22% within-person variability, TNF-α had 98.33% within-person variability, IL-12p70 had 99.01% within-person variability, IL-1β had 99.66% within-person variability, and IL-6 had 99.92% within-person variability. The correlation between protein ICCs and median intra-assay CVs was not significant (*r* = -.29, *p* = .421), suggesting that assay reliability did not drive these results.

**Table 4.** Reliability Results.

| Protein | Every-Three-Days Assessment Instability:<br>(% Within-person Variability) | Every-Six-Days Assessment Instability:<br>(% Within-person Variability) |
| --- | --- | --- |
| CRP | 54.18% | 44.99% |
| IL-10 | 79.26% | 89.75% |
| IL-4 | 94.96% | 96.98% |
| IFN-γ | 96.11% | 92.77% |
| IL-17A | 98.07% | 99.29% |
| IL-2 | 98.22% | 99.56% |
| TNF-α | 98.33% | 99.48% |
| IL-12p70 | 99.01% | 99.95% |
| IL-1β | 99.66% | 97.52% |
| IL-6 | 99.92% | 97.35% |

Note: CRP = C-reaction protein; IFN = interferon; IL = interleukin; TNF = tumor necrosis factor. Within-person variability = intraclass correlation coefficient converted to percentile and subtracted from 100%.

**Figure 1.**
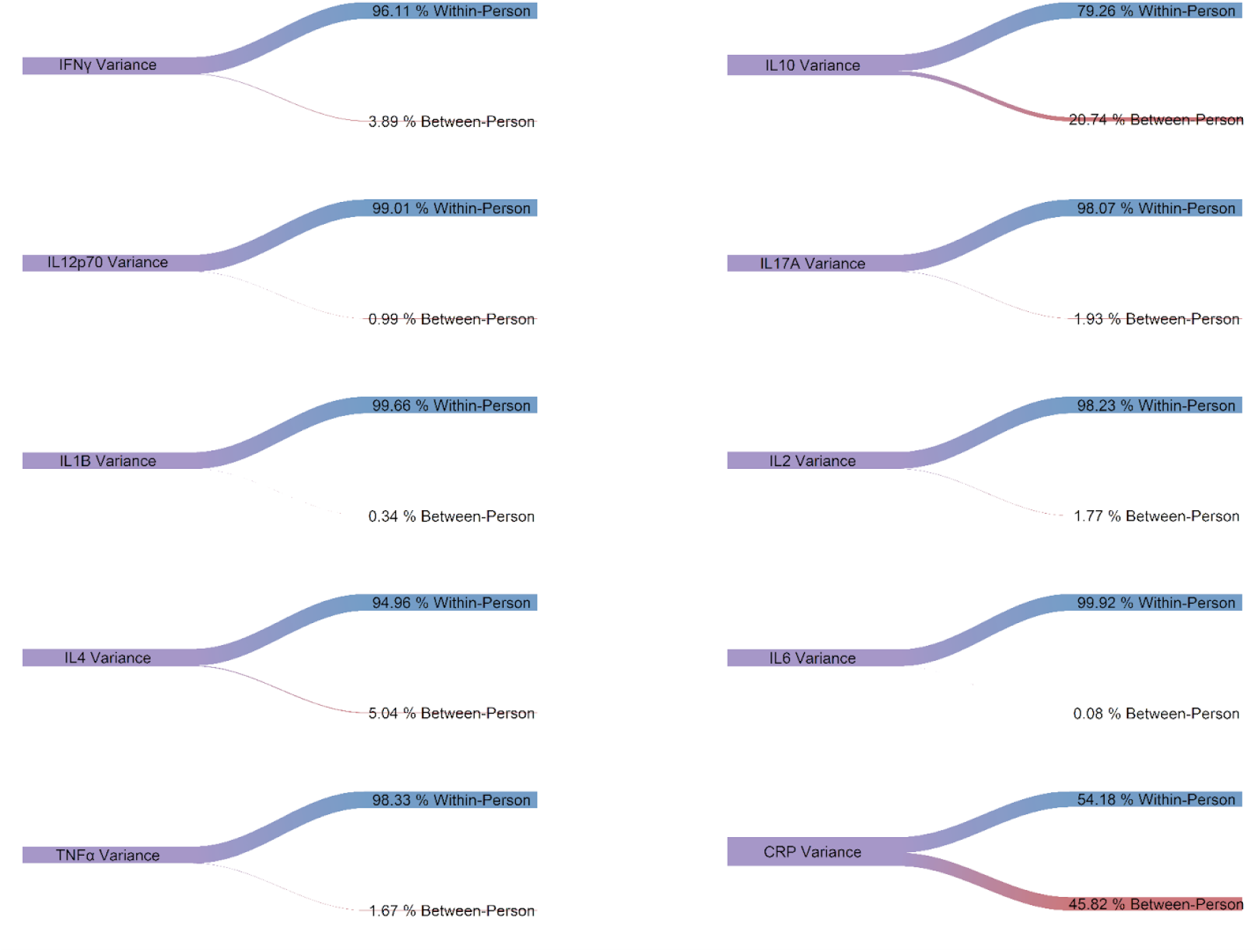
Riverplots of Between-vs. Within-Person Variability

To achieve an ICC of .5 (i.e., 50% between- and 50% within-person variability), CRP would need 9 assessments, IL-10 would need 31 assessments, IL-4 would need 151 assessments, IFN-γ would need 198 assessments, IL-17A would need 406 assessments, IL-2 would need 443 assessments, TNF-α would need 471 assessments, IL-12p70 would need 797 assessments, IL-1β would need 2,350 assessments, and IL-6 would need 10,058 assessments. As anticipated, reliability decreased for most proteins when reliability was tested using a 6-day assessment lag (see Table 4). Two of the four exceptions to this had over 99.5% within-person variability in the 3-day lag models, perhaps reflecting a ceiling effect.

## Discussion

Despite a great surge of interest in assessing inflammation in psychology and medicine^38,39^, the reliability of many peripheral immune protein levels remains unknown. To investigate, we combined a novel intensive longitudinal blood microsampling procedure and the transition to college as a socially salient, immersive period of stress and adjustment to study the temporal reliability of ten inflammatory proteins. The results suggest that, during times of stress and transition, the majority of variability in these inflammatory proteins are attributable to within-person changes instead of stable, between-person differences. None of the proteins assessed were reliable enough for their eight samples over 22 days to result in a profile characterized by at least 50% reliable, between-person differences. Conversely, all proteins were highly variable over time, with all proteins except C-reactive protein being characterized by over 95% within-person variability. Similar concerns emerged when considering average 3-day retest reliability, where even the highest correlation (.70) demonstrates that less than 50% of the variance in one assessment is associated with an assessment during the same 3 hour window just 3 days later. These results strongly support calls for greater investment in intensive longitudinal immune data collection in psychoneuroimmunology and immunopsychiatry research^18^, especially naturalistic studies investigating stress and/or social transitions.

These findings highlight concerns with longitudinal research focused on naturally occurring changes in these proteins as predictors or outcomes that have months- or years-long time lags between assessments. The degree of natural fluctuations observed here suggest that such designs do not have the temporal precision to capture change at the rate that it occurs. The greater the discrepancy between the rate of natural fluctuation and the not-frequent-enough immune assessment, the greater the risk for unmodeled time-varying confounders (e.g., substance use, exercise, diet); potential for the passing of time to dilute true, direct effects that operate on short time-scales; and risk for artificial amplification of effect sizes due to positive feedback loops over time.

In addition to better complimenting the natural temporal reliability of these inflammatory proteins, intensive longitudinal designs offer additional benefits for the clinical translatability of research^40^. Many researchers study questions about how change in inflammation is associated with changes in other health processes (and vice versa). Repeated measures data facilitates analyses that isolate within-person variance, which is of greater causal relevance than observational research focused strictly on between-person variability (e.g., baseline inflammation predicting future depression) or that fails to disaggregate within-from between-person variability^41^. Further, power for tests of within-person effects is greatly influenced by the number of observations per person. As a hypothetical example, if a hypothesis needs 10 observations per person to be powered, collecting those assessments over weeks or months will reduce some of the financial and logistical hassle of maintaining relationships with research participants and lab infrastructure compared to 5 years of biannual assessments. These results also provide estimates for 3-day retest reliability and intra-class correlation coefficients that can be used for power-analyses for other longitudinal research with these proteins.

### Strengths and Limitations

This study features several key strengths for investigating the temporal reliability of proinflammatory proteins during a critical period of stress and social transition. First, the high intensity data collection of blood based inflammatory proteins across multiple participants is uniquely synergistic for this research question compared to typical longitudinal studies with months-to-years long time lags^42^. Second, the focus on blood-based proteins helps address the lack of physiometric research with this methodology, as many physiometric studies in this area focus on salivary or urinary immune assessments^23,27,31^. Third, while some intensive longitudinal immune assessments have been published as part of case studies, these preclude the ability to compare between-vs. within-person variance due to the focus on a single individual^43^. Fourth, assaying the cytokines in triplicate ameliorates some concerns about multiplex reliability by (a) allowing to average across three assessments instead of two and (b) facilitating the identification and removal of individual assays that were extreme outliers of questionable validity without reducing sample size. Fifth, executing this study during a period of stress and transition complements existing focus on how stress modulates the immune system in ways that may be relevant for health. Sixth, collecting time-verified immune assessments in the morning helped to prevent immune confounding due to diurnal variations or differences in biobehavioral patterns that emerge across the day.

The results must also be considered in light of several limitations. First, like all measurement research, it is unclear the extent to which these estimates will generalize to other contexts (e.g., during periods of relative stability and relaxation) or samples (e.g., medical, older adult). Second, the three-day measurement lag over 22 days was arbitrarily chosen given the lack of previous research into these properties and financial limits of the grant used to fund this work. Additional work needs to be done with both shorter and longer time lags. Similarly, the panel of proteins selected were chosen to cover a breadth of pro- and anti-inflammatory processes; however, this is far from a fully comprehensive assessment of immune biology.

## Conclusion

In conclusion, the present data underscore that, during times of stress and transition, the majority of variability in these inflammatory proteins are attributable to within-person changes instead of stable, between-person differences. As such, studies investigating the interplay between these inflammatory proteins and other health processes, especially in the context of stress, would benefit from high frequency assessment of inflammatory biology^18^. This shift in data collection strategy, compared to the status-quo of months-to-years long time lags, would better capture inflammatory change as it occurs and complement the within-person effects that are central to the mechanistic hypotheses that are central to the clinical relevance of the field.

